# Identification of Characteristic Genomic Markers in Human Hepatoma Huh7 and Huh7.5.1-8 Cell Lines

**DOI:** 10.1101/2020.02.17.953281

**Authors:** Masaki Kawamoto, Toshiyuki Yamaji, Kyoko Saito, Kazuhiro Satomura, Toshinori Endo, Masayoshi Fukasawa, Kentaro Hanada, Naoki Osada

**Affiliations:** Graduate School of Information Science and Technology, Hokkaido University, Sapporo 060-0814, Japan; Department of Biochemistry & Cell Biology, National Institute of Infectious Diseases, Tokyo 162-8640, Japan; Faculty of Information Science and Technology, Hokkaido University, Sapporo 060-0814, Japan; Global Station for Big Data and Cybersecurity, GI-CoRE, Hokkaido University, Sapporo 060-0814, Japan

**Keywords:** Huh7, genome sequencing, cell lines, hepatitis C virus

## Abstract

The human hepatoma-derived Huh7 cell line and its derivatives (Huh7.5 and Huh7.5.1) have been widely used as a convenient experimental substitute for primary hepatocytes. In particular, these cell lines represent host cells suitable for propagating the hepatitis C virus (HCV) *in vitro*. The Huh7.5.1-8 cell line, a subline of Huh7.5.1, can propagate HCV more efficiently than its parental cells. To provide genomic information for cells’ quality control, we performed whole-genome sequencing of Huh7 and Huh7.5.1-8 and identified their characteristic genomic deletions, some of which are applicable to an in-house test for cell authentication. Among the genes related to HCV infection and replication, 53 genes were found to carry missense or loss-of-function mutations likely specific to the Huh7 and/or Huh7.5.1-8. Eight genes, including *DDX58* (*RIG-I*), *BAX, EP300*, and *SPP1* (osteopontin), contained mutations observed only in Huh7.5.1-8 or mutations with higher frequency in Huh7.5.1-8. These mutations might be relevant to phenotypic differences between the two cell lines and may also serve as genetic markers to distinguish Huh7.5.1-8 cells from the ancestral Huh7 cells.

## Introduction

Quality control of cells is crucial for biopharmaceutical manufacturing as well as research. In research, many cell lines have been misidentified (http://iclac.org/wp-content/uploads/Cross-Contaminations-v8_0.pdf), and various academic journals have begun requesting proper authentication of cells used in studies. Cell lines can be identified by genotyping multiple common genetic variants, such as microsatellite markers and single nucleotide variants (SNVs)(Gilbert et al., 1990, Ghandi, 2019 #11972). However, such common markers would not be useful for discriminating between cell lines that originated from the same ancestral cell line. Thus, rational, easy in-house methods of identifying cell lines are widely desired. In particular, the mutation rate of deletion is much lower than that of point mutations(Wang et al., 2008). Cell line-specific deletions are applicable to cell line authentication of African green monkey kidney-derived Vero cells (Osada et al., 2014),(Sakuma et al., 2018), which are widely used in virologic research and human vaccine production.

HuH-7 (hereafter Huh7) is a permanent cell line established from male hepatoma tissue, which was surgically removed from a 57-year-old Japanese male in 1982(Nakabayashi et al., 1982). Huh7 and its derivatives have been used as a convenient experimental substitute for primary hepatocytes. Approximately 80% of hepatocellular carcinoma incidents in humans are caused by hepatitis viruses; in all hepatocellular carcinoma incidents, ∼25% were caused by hepatitis C virus (HCV) and ∼53% were caused by hepatitis B virus (HBV)(Perz et al., 2006). HCV is a positive-stranded RNA virus that infects humans, causing acute and chronic liver diseases. The chronic infection eventually leads to severe symptoms, such as hepatitis and hepatocellular carcinoma. Thus, permanent cell lines suitable for investigating human hepatitis viruses have always been invaluable. After the establishment of Huh7, several studies have attempted to derive cell lines that are more permissive to HCV than Huh7 (Blight et al., 2002; Feigelstock et al., 2010). The whole-genome sequence of Huh7 was determined in a large-scale genome sequencing project of various cancer cell lines (Ghandi et al., 2019); however, a detailed study of the Huh7 genome sequence has not been performed. Huh7.5, a subline of Huh7, was established as a highly permissive cell line to replicate subgenomic and genomic HCV RNA in 2002 (Blight et al., 2002). Interestingly, Huh7.5, but not the ancestral Huh7, has a missense mutation in *DDX58* (or *RIG-I*) gene, which participates in intracellular antiviral defense (Sumpter et al., 2005). Although cell culture systems that recapitulate the HCV life cycle have not been developed for a long time, the JFH-1 HCV strain has been found to produce infective progenitor virions after introduction to Huh7.5 cells(Wakita et al., 2005). Huh7.5.1, a subline of Huh7.5 cells, was subsequently established as a cell line with the intent to generate a cell line that cultures JFH-1 more efficiently than Huh7.5 (Zhong et al., 2005). Although Huh7.5.1 displayed faster viral replication kinetics than Huh7, Huh7 and Huh7.5.1 eventually produced a similar level of viral titers (Zhong et al., 2005). We previously found that the expression level of CD81, which is a plasma membrane protein essential for HCV infection (Petracca et al., 2000), was not uniform in the Huh7.5.1 cells, and we thus obtained Huh7.5.1-8, a subclone of Huh7.5.1, in which CD81 is stably expressed on cell surfaces (Shirasago et al., 2015). Huh7.5.1-8 cells exhibit ∼10-fold greater permissiveness to HCV than Huh7.5.1 cells (Shirasago et al., 2015). HCV culture systems have provided robust assay systems to screen anti-HCV compounds, resulting in the development of various anti-HCV medicines that are currently marketed, although anti-HCV vaccines have not been developed.

We determined the whole-genome sequences of Huh7 and Huh7.5.1-8 cell lines and found characteristic genomic deletions in Huh7 and Huh7.5.1-8, some of which are applicable to an in-house test for cell authentication. In addition, among the genes related to HCV infection and replication, 53 genes were found to carry missense or loss-of-function (LoF) mutations that were not registered in the public germline mutation databases, but were found in the Huh7 and/or Huh7.5.1-8. Among them, 8 genes, including *DDX58* (*RIG-I*), *BAX, EP300*, and *SPP1* (osteopontin), contained mutations observed only in Huh7.5.1-8 or mutations with higher frequency in Huh7.5.1-8.

## Materials and Methods

### DNA sample preparation and sequencing

DNA samples of Huh7 hepatocarcinoma cell line (JCRB0403) and Huh7.5.1-8 cell line were obtained from the Japanese Collection of Research Bioresources (JCRB) Cell Bank. We confirmed that the HCV replication is slower in JCRB0403 than Huh7.5.1-8 (data not shown). For both samples, fragment libraries (average fragment sizes of ∼560 bp) were constructed using TruSeq DNA PCR-Free LT Library Prep Kit (Illumina, San Diego, CA, USA). Paired-end sequences of 150 bp long were determined using HiSeq X (Illumina). Approximately 1.8 billion sequence reads were obtained from each sample. We examined whether viral sequences of HBV and HCV are integrated into the two genomes using VirusFinder 2 software(Wang et al., 2015). The sequences were deposited in the DDBJ DRA database under project ID PRJDB7928.

### RNA-seq data

We retrieved previously obtained RNA-seq data from Huh7.5.1 and Huh7.5.1-8 (DRR018792 and DRR018793, respectively). Because the initial quality check for the RNA-seq data showed some of the read bases had relatively low quality, we applied Trimmomatic software (version 0.36)(Bolger et al., 2014) to filter out low-quality reads. After trimming low-quality bases of average quality score <20 (window size: 4), reads shorter than 75 bp were filtered out. RNA-seq reads were mapped to the reference human genome using HISAT2 (Kim et al., 2015).

### SNV calling

Genomic reads were mapped to the reference human genome sequence (GRCh38 primary assembly downloaded from the Ensembl database) using the BWA MEM algorithm (version 0.7.15-r1140)(Li and Durbin, 2009) with a default parameter setting. The read mapping rates of Huh7 and Huh7.5.1-8 were 99.88% and 99.87%, respectively. To generate alignment files for variant calling, GATK Best Practices Pipeline 3.0, which includes duplicated read filtering, realignment around indels, and recalibration of base quality score, was applied to the initial alignments(McKenna et al., 2010).

The sites with read depths of ≥14 and ≤100 were used for the following variant calling. The number of sites with ≥14 read depth covered >95% of the reference human genome in our dataset. Initial SNV calling was performed using VarScan (version 2.4.3) (Koboldt et al., 2012) with a base quality score cutoff of 15 and a variant allele frequency cutoff of 0.1%. Multiallelic SNVs were removed from the analytical pipeline. Known germline SNVs registered in the public database (dbSNP_149) were filtered out using the SelectVariants program in the GATK package(McKenna et al., 2010). However, when a variant nucleotide in Huh7 and/or Huh7.5.1-8 was different from that in the dbSNP, those variants were kept for further analyses as a novel variant.

We defined three categories of newly identified SNVs according to their frequency in Huh7 and Huh7.5.1-8. We tested whether the SNV frequency is higher in Huh7 or Huh7.5.1-8 with statistical significance. Statistical significance was evaluated using the χ^2^ test and a false discovery rate of 0.05 (Benjamini and Hochberg, 1995). The SNVs with higher frequency in Huh7 and Huh7.5.1-8 were categorized into “Huh7-predominant” and “Huh7.5.1-8-predominant” categories, respectively. The SNVs that did not pass the criteria were further classified into “shared” SNVs when the variant frequencies in both samples exceeded 0.25.

### Structural variant calling

Indels shorter than 50 bp were identified, using VarScan, with the same pipeline for SNV calling. Indels that exactly matched to known germline indels (Mills and 1000G gold standard indels (Mills et al., 2006)) were filtered out from further analyses. Accordingly, short indels were classified into three categories: Huh7-predominant, Huh7.5.1-8-predominant, and shared short indels.

Long indels (≥50 bp) were identified using Manta (version 1.1.0) (Chen et al., 2016). As with the SNV calling, we only considered the sites with read depths of ≥14 and ≤100. The estimated structural variant frequency in Manta was difficult when we did not obtain a sufficient number of reads spanning breakpoints. Therefore, we classified the structural variants into the above three categories without considering variant frequencies. Large insertions that match to the insertions in Mills and 1000G gold standard dataset and large deletions that overlapped (≥50% length of the estimated deletion size in the cell lines) with the deletions of Mills and 1000G gold standard and/or 1000 Genome Phase 3 structural variants (downloaded from ftp://ftp.1000genomes.ebi.ac.uk/vol1/ftp/phase3/) were filtered out.

### Functional annotation

Functional annotation of mutations was performed using the snpEff software (Cingolani et al., 2012) with an annotation data snpEff_v4_3_GRCh38.86. Using the annotation information, SNVs with strong phenotypic effects (missense mutations, nonsense mutations, and mutations at splicing acceptor-donor sites) were extracted. Functional annotation of genes was performed using the DAVID bioinformatics resource(Jiao et al., 2012). We used the following five fields in DAVID output: disease, functional categories, gene ontology, pathways, and protein domains. The protein-protein association networks were retrieved from the STRING database (Szklarczyk et al., 2015). The Python scripts used in this study were deposited in GitHub (https://github.com/J1011Ibb14029/kwmtools).

### Sequence conservation

We investigated the evolutionary conservation at mutated amino acid sites using the amino acid sequences of 12 nonhuman vertebrate species: *Pan troglodytes, Pan paniscus, Gorilla gorilla, Pongo abelii, Macaca mulatta, Cricetulus griseus, Felis catus, Desmodus rotundus, Phascolarctos cinereus*, and *Gallus gallus.* Sequence data were retrieved and aligned using MEGAX (Knyaz et al., 2018), and sequence alignment was performed with the MUSCLE algorithm (Edgar, 2004).

### Experimental validation of deletions

The HeLa cervical carcinoma cell line (JCRB9004) was obtained from the JCRB Cell Bank and used as a control. Genome DNA was prepared from the cell lines using the Blood Genomic DNA Extraction Mini Kit (Favorgen, Ping-Tung, Taiwan, ROC). Deletions in the genomes of Huh7 and Huh7.5.1-8 cells were verified by PCR experiments using a previously reported procedure (Sakuma et al., 2017). Briefly, PrimeSTAR GXL DNA Polymerase (Takara Bio, Otsu, Japan) was used for amplification. The reaction mixture, containing 60 ng genome DNA, was denatured at 98°C for 1 min and then subjected to 40 cycles, consisting of 98°C for 10 s, 61°C for 15 s, and 68°C, for 1 min. The amplified products were electrophoresed on an agarose gel and visualized with the Gel Doc EZ imager (Bio-Rad, Hercules, CA, USA). Then, 1 kb Plus DNA Ladder was used as a molecular marker (Thermo Fisher Scientific, Waltham, MA, USA).

### Sanger sequencing of DDX58 gene

Genomic DNA was prepared from 1 × 10^6^ cells using the Blood Genomic DNA Extraction Mini Kit (Favorgen) and used as a template to amplify a 446 bp fragment corresponding to bases 30,196 to 30,641 of the human DDX58 genic region (National Center for Biotechnology Information accession number NG_046918.1; 78,023 bp) using PCR. PCR was performed with 35 cycles of amplification (95°C for 10 s, 57°C for 5 s, and 72°C for 5 s) on the PCR Thermal Cycler Dice (Takara Bio) using PrimeSTAR MAX DNA polymerase (Takara Bio), forward primer 5′-GTGGCTTGGTGAAGAATGGGCACAG-3′ (bases 30,196–30,220), and reverse primer 5′-CTCAGACTAAGAGGCATGAACTATAAGTGG-3′ (complementary to bases 30,612–30,641). The resulting fragments were separated in an agarose gel, purified using with a gel/PCR extraction kit (FastGene; Nippon Genetics, Tokyo, Japan), and directly sequenced by the Sanger method using forward primer 5′-CCCTATTTGGGAAGGTCTGGTGATC-3′ (bases 30,291–30,315) and reverse primer 5′-CACTTTTACAGTATTGTCAAGCAGC-3′ (complementary to bases 30,564–30,588) by Eurofins Genomics K.K. (Tokyo, Japan).

## Results and Discussion

### Identification of mutations in Huh7 lineages

We obtained 41.0- and 41.1-fold coverages of whole-genome sequence reads from Huh7 (JCRB0403) and Huh7.5.1-8 cells, respectively. No sequences related to HBV and HCV were detected in the genomes using the VirusFinder2 pipeline, as reported previously for HBV using *in situ* hybridization (Tay et al., 1990). To identify novel mutations, we filtered out mutations matched to the previously known germline SNVs and indels from further analyses (see Materials and Methods). These mutations were not found in currently available human mutation database, suggesting that most (even if not all) of these newly-identified SNVs specifically occurred during the establishment of the Huh7 cell line and/or the development of the liver cancer in the patient, from which the cell line was established. For the following analysis, we further classified mutations into three categories: 1) Huh7-predominant mutations, of which frequency is higher in Huh7 than in Huh7.5.1-8, 2) Huh7.5.1-8-predominant mutations, of which mutation frequency is higher in Huh7.5.1-8 than in Huh7, and 3) shared mutations, of which frequency is almost equal between the two cell lines. The SNVs of higher frequency in Huh7 than in Huh7.5.1-8, with statistical significance (false discovery rate of 0.05), were classified as Huh7-predominant SNVs and vice versa. The SNVs that did not pass the criteria were further classified into shared SNVs when the variant frequencies in both samples exceeded 0.25. Table 1 summarizes the number of mutations, including SNVs, insertions, and deletions.

**Table 1.** Summary of newly identified variants in Huh7 and Huh7.5.1-8.

| Mutation type | Huh7-predominant | Huh7.5.1-8-predominant | Shared |
| --- | --- | --- | --- |
| SNV | 4,094 | 39,497 | 183,856 |
| Insertion | 911 | 933 | 27,206 |
| Deletion | 1,898 | 8,948 | 126,084 |

As expected, many new mutations had higher frequencies in Huh7.5.1-8 than in Huh7 (Huh7.5.1-8-predominant class), because Huh7.5.1-8 should have experienced a larger number of cell passages. However, Huh7 also had a substantial number of mutations that were not found in Huh7.5.1-8. The observation was, presumably, accounted for by the fact that the seed stock of Huh7 cells we sequenced was not a direct ancestral cell seed that was used to establish Huh7.5. In total, we identified 394,568 new mutations, including 227,447 SNVs and 29,050 insertions, and 136,930 deletions.

Most of the mutations were shared between the two cell lines. These shared mutations were most likely to have arisen before the establishment of the Huh7 cell line. Heterogeneity of mutations is often seen not only among different cancer types but also among cancer cells of different patients diagnosed with the same cancer type. Indeed, the amount and pattern of new mutations in liver cancer tissues were shown to be considerably diverse (Fujimoto et al., 2015). In our study, Huh7 and Huh7.5.1-8 contained a relatively large number of new mutations, although it is impossible to obtain germline cells of the patient from whom Huh7 derived to sequence their genome, and some of the mutations might have occurred in the germline cells. Notably, the number of insertions and deletions was considerably high, and the ratio of indels to SNVs was much higher than the ratio observed in germline cells (Wang et al., 2008). Our validation, using polymerase chain reaction (PCR), confirmed that all six amplified regions contained specific deletions in the cell line DNA, indicating that the false-positive rate would not be substantially high in our variant calling pipeline.

Interestingly, we identified one nonsynonymous mutation (K70R) in the *POLD3* gene, which plays an important role in high-fidelity DNA replication (Johansson and MacNeill, 2010) in both Huh7 and Huh7.5.1-8. In all 12 nonhuman vertebrates we analyzed, POLD3 had lysine residues at site 70, indicating that this amino acid site has a very important function in the protein. *POLD3* mutations have been identified repeatedly in cancer cells (Wang et al., 2014), and they contribute to chromosomal instability (Minocherhomji et al., 2015). We suspected that K70R mutations in POLD3 partly explain the large number of SNVs and indels and the frequent chromosome copy number changes in Huh7(Kasai et al., 2018).

### Deletion markers for Huh7 cell lines

To establish efficient deletion markers characteristic of Huh7 cells, we designed PCR primer sets that test the presence of deletion in genomic DNA using the dataset of homozygous deletions identified in whole-genome sequencing. Four long deletions, ranging from 7,400 to 262,700 bp in length, and four short deletions, ranging from 645 to 1018 bp in length, were targeted for PCR amplification. We designated regions harboring long and short deletions as DL and DS regions, respectively. In our genome sequencing analysis, Five indels were expected to be present only in Huh7.5.1-8, whereas the others were expected to be shared between Huh7 and Huh7.5.1-8. We used a DNA sample from HeLa cells as a control. Primer sequences and detailed information for each region are presented in Supplementary Table 1.

We successfully amplified six regions by genomic PCR and confirmed that they all harbored deletions of expected sizes in Huh7 and/or Huh7.5.1-8, but did not show the signature deletion in the HeLa cells (Figure 1). In total, four deletions (DL1–3 and DS2) were present only in Huh7 and Huh7.5.1-8, and two deletions (DS1 and DS3) were observed only in Huh7.5.1-8. The results are summarized in Supplementary Table 2.

**Figure 1.**
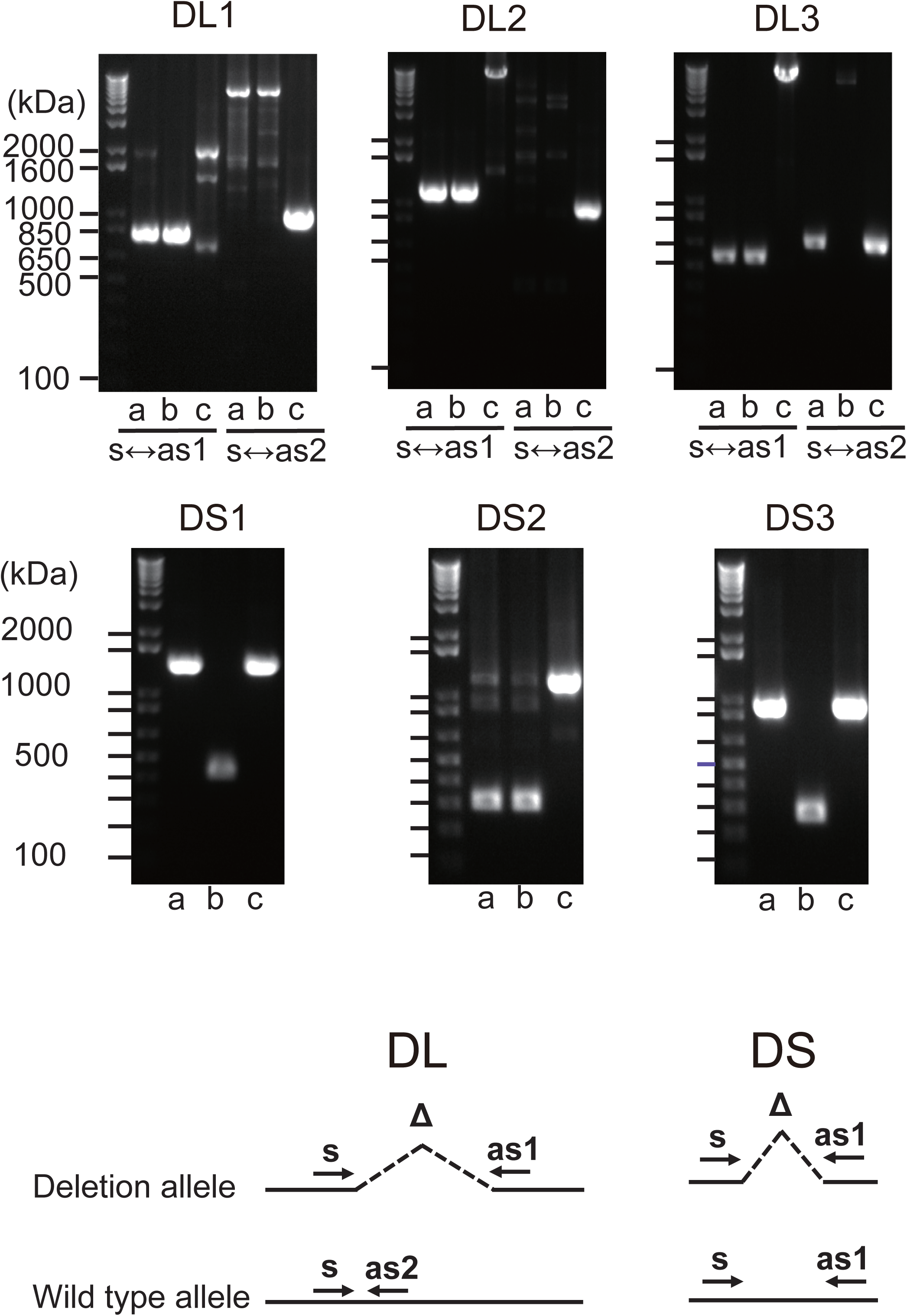
Evaluation of deletions identified in Huh7 and Huh7.5.1-8 using PCR. Samples in lanes a to c are DNA from Huh7, Huh7.5.1-8, and HeLa cells, respectively. Size markers are indicated by horizontal lines. The designs for PCR amplification and expected results are shown on the bottom right. The region name is given for each gel image. Prefixes DL and DS represent whether the expected deletion length is long (>7.4 kbp) or short (<0.9 kbp). For DL regions, primer pairs used for the experiments are shown below each image. For DS regions, primer pairs s and as1 were used for all regions. Each gel image was cropped from the full-sized gels, but different lanes were not merged into a single image.

### Classification of newly identified SNVs

The newly identified SNVs were further classified into missense mutations, nonsense mutations, mutations at splicing signal sites, and other mutations. In addition, frameshift and large indels harboring protein-coding sequences (including gene fusions) were classified as LoF mutations. Nonsense mutations and mutations at splicing signal sites were also classified as LoF mutations.

Among these mutations, we selected genes that have missense and/or LoF mutations. We confirmed that missense mutations are present in the RNA-seq read data, which means that mutated alleles are actually expressed in the cells. In total, 255 and 431 genes were found to have missense and LoF mutations, respectively. The complete list of genes is shown in Supplementary Data 1.

We narrowed down the gene list and chose several genes that might be relevant to the HCV replication process. We reviewed previous research and selected keywords that are related to HCV infection and replication (Scheel and Rice, 2013; Bukh, 2016): autophagy, apoptosis, antiviral defense, hepatitis C, innate immune response, and serine protease. In addition, we surveyed the genes involved in protein-protein association networks with nine core genes that showed a strong influence on HCV infection and replication (*CD36, CD81, CLDN1, EGFR, EPHA2, LDLR, OCLN, PPIA*, and *SCARB1*).

We identified 53 candidate genes, including 12 autophagy-related genes, 22 apoptosis-related genes, 3 antiviral defense genes, 14 HCV-related genes, 8 innate immune-related genes, 4 serine proteases, and 4 genes involved in the HCV core-gene network. Eleven genes were categorized into two different classes. The list of genes and mutations is summarized in Table 2. Two HLA genes, *HLA-DRB1* and *HLA-DRB5*, related to an acquired immune system, contained several LoF mutations shared between Huh7 and Huh7.5.1-8, albeit those genes were not included in the 53 genes in Table 2 since cultured cells lack the acquired immune system. Interestingly, among the 53 candidate genes, 8 genes contained Huh7.5.8-1+ mutations, but none carried Huh7-predominant mutations. Among the 8 genes, 4 genes (*BAX, COL6A3, DEFB104B*, and *SIRPB1*) had LoF mutations, whereas the other 4 genes (*DDX58, EP300, SPP1*, and *ZNF654*) had missense mutations (Table 2). Future studies will be required to elucidate whether these mutations are relevant to the phenotypes of the Huh7 cell lineage.

**Table 2.**
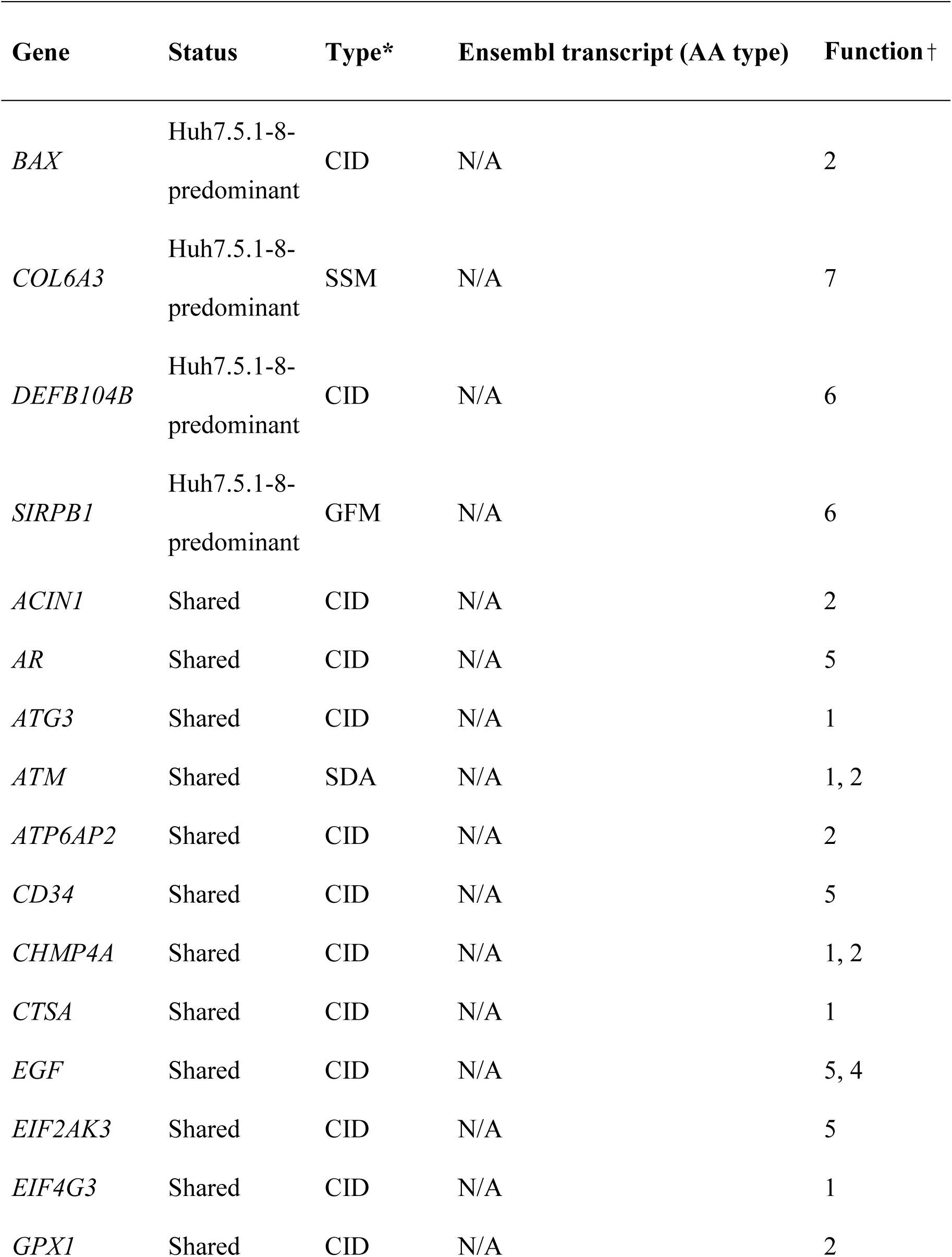
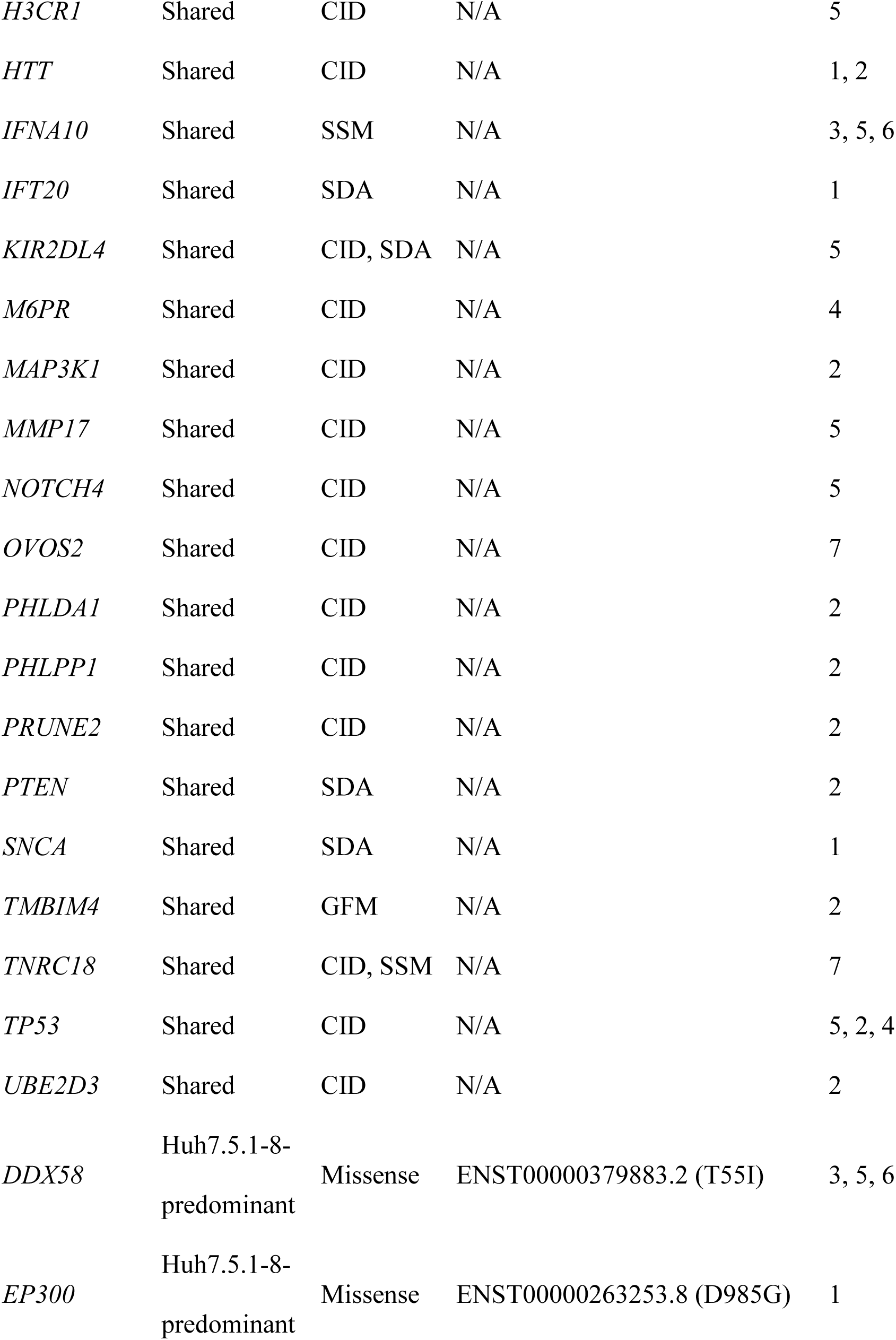
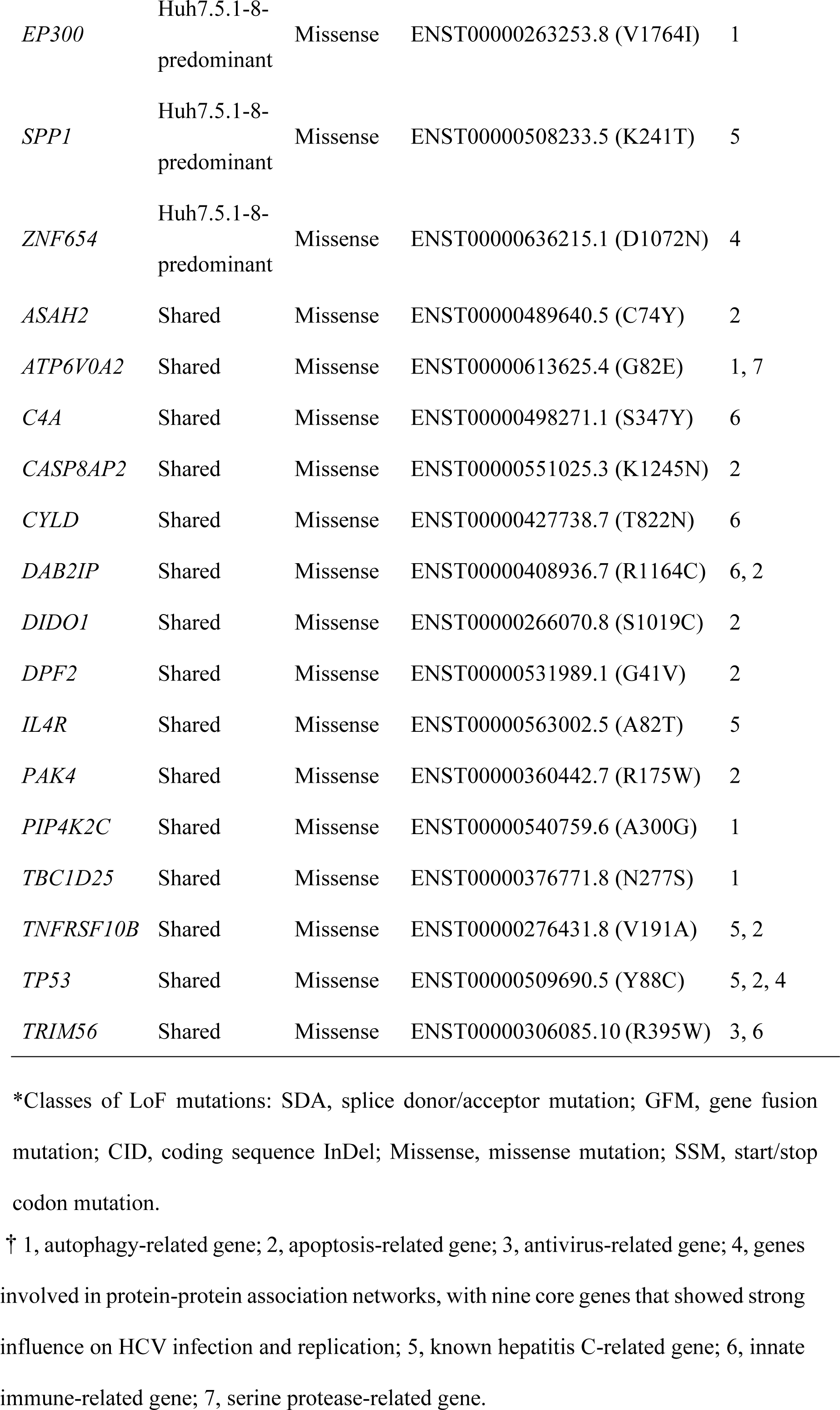
Mutations potentially contributing HCV permissiveness to Huh7 and Huh7.5.1-8.

| Gene | Status | Type* | Ensembl transcript (AA type) | Function† |
| --- | --- | --- | --- | --- |
| *BAX* | Huh7.5.1-8-predominant | CID | N/A | 2 |
| *COL6A3* | Huh7.5.1-8-predominant | SSM | N/A | 7 |
| *DEFB104B* | Huh7.5.1-8-predominant | CID | N/A | 6 |
| *SIRPB1* | Huh7.5.1-8-predominant | GFM | N/A | 6 |
| *ACIN1* | Shared | CID | N/A | 2 |
| *AR* | Shared | CID | N/A | 5 |
| *ATG3* | Shared | CID | N/A | 1 |
| *ATM* | Shared | SDA | N/A | 1, 2 |
| *ATP6AP2* | Shared | CID | N/A | 2 |
| *CD34* | Shared | CID | N/A | 5 |
| *CHMP4A* | Shared | CID | N/A | 1, 2 |
| *CTSA* | Shared | CID | N/A | 1 |
| *EGF* | Shared | CID | N/A | 5, 4 |
| *EIF2AK3* | Shared | CID | N/A | 5 |
| *EIF4G3* | Shared | CID | N/A | 1 |
| *GPX1* | Shared | CID | N/A | 2 |
| *H3CR1* | Shared | CID | N/A | 5 |
| *HTT* | Shared | CID | N/A | 1, 2 |
| *IFNA10* | Shared | SSM | N/A | 3, 5, 6 |
| *IFT20* | Shared | SDA | N/A | 1 |
| *KIR2DL4* | Shared | CID, SDA | N/A | 5 |
| *M6PR* | Shared | CID | N/A | 4 |
| *MAP3K1* | Shared | CID | N/A | 2 |
| *MMP17* | Shared | CID | N/A | 5 |
| *NOTCH4* | Shared | CID | N/A | 5 |
| *OVOS2* | Shared | CID | N/A | 7 |
| *PHLDA1* | Shared | CID | N/A | 2 |
| *PHLPP1* | Shared | CID | N/A | 2 |
| *PRUNE2* | Shared | CID | N/A | 2 |
| *PTEN* | Shared | SDA | N/A | 2 |
| *SNCA* | Shared | SDA | N/A | 1 |
| *TMBIM4* | Shared | GFM | N/A | 2 |
| *TNRC18* | Shared | CID, SSM | N/A | 7 |
| *TP53* | Shared | CID | N/A | 5, 2, 4 |
| *UBE2D3* | Shared | CID | N/A | 2 |
| *DDX58* | Huh7.5.1-8-predominant | Missense | ENST00000379883.2 (T55I) | 3, 5, 6 |
| *EP300* | Huh7.5.1-8-predominant | Missense | ENST00000263253.8 (D985G) | 1 |
| *EP300* | Huh7.5.1-8-predominant | Missense | ENST00000263253.8 (V1764I) | 1 |
| *SPP1* | Huh7.5.1-8-predominant | Missense | ENST00000508233.5 (K241T) | 5 |
| *ZNF654* | Huh7.5.1-8-predominant | Missense | ENST00000636215.1 (D1072N) | 4 |
| *ASAH2* | Shared | Missense | ENST00000489640.5 (C74Y) | 2 |
| *ATP6V0A2* | Shared | Missense | ENST00000613625.4 (G82E) | 1, 7 |
| *C4A* | Shared | Missense | ENST00000498271.1 (S347Y) | 6 |
| *CASP8AP2* | Shared | Missense | ENST00000551025.3 (K1245N) | 2 |
| *CYLD* | Shared | Missense | ENST00000427738.7 (T822N) | 6 |
| *DAB2IP* | Shared | Missense | ENST00000408936.7 (R1164C) | 6, 2 |
| *DIDO1* | Shared | Missense | ENST00000266070.8 (S1019C) | 2 |
| *DPF2* | Shared | Missense | ENST00000531989.1 (G41V) | 2 |
| *IL4R* | Shared | Missense | ENST00000563002.5 (A82T) | 5 |
| *PAK4* | Shared | Missense | ENST00000360442.7 (R175W) | 2 |
| *PIP4K2C* | Shared | Missense | ENST00000540759.6 (A300G) | 1 |
| *TBC1D25* | Shared | Missense | ENST00000376771.8 (N277S) | 1 |
| *TNFRSF10B* | Shared | Missense | ENST00000276431.8 (V191A) | 5, 2 |
| *TP53* | Shared | Missense | ENST00000509690.5 (Y88C) | 5, 2, 4 |
| *TRIM56* | Shared | Missense | ENST00000306085.10 (R395W) | 3, 6 |

\*Classes of LoF mutations: SDA, splice donor/acceptor mutation; GFM, gene fusion mutation; CID, coding sequence InDel; Missense, missense mutation; SSM, start/stop codon mutation.
† 1, autophagy-related gene; 2, apoptosis-related gene; 3, antiviral-related gene; 4, genes involved in protein-protein association networks, with nine core genes that showed strong influence on HCV infection and replication; 5, known hepatitis C-related gene; 6, innate immune-related gene; 7, serine protease-related gene.

One of the genes, *DDX58*, which is also known as *RIG-I*, encodes an RNA helicase and has an important function for innate antiviral response (Yoneyama et al., 2004). The mutation is, therefore, a strong candidate for the high permissiveness of HCV in the Huh7.5 lineage. It is noteworthy that our whole-genome sequence approach also identified this gene. However, we found a difference in the mutation frequency between the results of this and previous studies. This incongruence is discussed in the next subsection.

Because we do not have sufficient space to discuss all 53 candidate genes for the higher permissiveness of HCV in detail, we here selected three key genes that have important viral replication functions.

BAX, which has an LoF mutation observed only in Huh7.5.1-8, forms a dimer with BCL2 and activates apoptosis (Oltval et al., 1993). In the Huh7 cell lineage, the *TP53* gene, which encodes P53 protein and controls the apoptotic process through BAX (Toshiyuki and Reed, 1995), has a frameshift deletion (Table 2). Therefore, BAX may positively regulate apoptosis in Huh7. In contrast, in Huh7.5.1-8, 22% of *BAX* alleles carried frameshift insertions, which would decrease *BAX*’s overall activity. The apoptotic activity, therefore, might have decreased in Huh7.5.1-8 compared with Huh7, which might result in HCV’s higher replication efficiency.

Another candidate gene, *EP300*, which encodes adenovirus early region 1A-associated protein p300 (Eckner et al., 1994; Ogryzko et al., 1996), harbored two missense mutations (D985G and V1764I) in Huh7.5.1-8. *EP300* has the function of suppressing the autophagy process(Pietrocola et al., 2014). Previous studies have shown that HCV and other flaviviruses replicate in host cells using autophagosomes (Levine et al., 2011; Fahmy and Labonté, 2017). We found that the valine residue at site 1764 was highly conserved among vertebrates. The mutation may have reduced *P300*’s function for autophagy suppression and contributed to efficient HCV replication.

The last gene we present here is *SPP1* (osteopontin), which is involved in the remodeling process of bones (Kahles et al., 2014). Previous studies found that (1) the gene expression of *SPP1* in liver tissue was increased according to the progress of hepatitis C (Asselah et al., 2005), (2) *SPP1* enhanced autophagy in human hepatocellular carcinoma cells (Liu et al., 2016), and (3) *SPP1* interacted with HCV proteins and helped replicate and assemble HCV(Iqbal et al., 2018). Indeed, it was reported that *SPP1* expression is up-regulated in Huh7.5 compared with Huh7 (Choi et al., 2014). Our study found that 46% of *SPP1* alleles in Huh7.5.1-8 carried missense mutations of K241T, whereas the frequency of mutation in Huh7 was 0%. The frequency of mutations in both Huh7.5.1 and Huh7.5.1-8 transcripts was close to 50%, indicating the K241T mutation had already been acquired in Huh7.5.1, and the expression of the transcript with threonine allele would not be up-regulated by HCV infection in an allele-specific manner. The missense mutation, as well as the overall elevation of gene expression, may have increased the replication efficiency of the HCV in Huh7 derivatives.

### Missense mutation in RIG-I (T55I) was heterologous in the Huh7.5.1-8 genome

As described in the above subsection, we found a T55I mutation in RIG-I that could contribute to the high permissiveness to HCV in Huh7.5.1-8. However, both genome sequencing and RNA-seq data showed the mutation is heterologous in Huh7.5.1-8, contradicting previous findings that the mutation is homozygous in Huh7.5 (Sumpter et al., 2005). Therefore, we resequenced the genomic DNA from Huh7 and Huh7.5.1-8 using a Sanger sequencer and verified that the mutation is absent in Huh7 but heterologous in Huh7.5.1-8 (Figure 2). Our recent study showed that a 55T allele transcript harbors an allele-specific large deletion and, presumably, nonfunctional in Huh7.5.1-8 (Saito et al., unpublished results). The results showed that, although T55I mutation is heterozygous in the Huh7.5.1-8 genome, all full-length RIG-I proteins have the T55I mutation.

**Figure 2.**
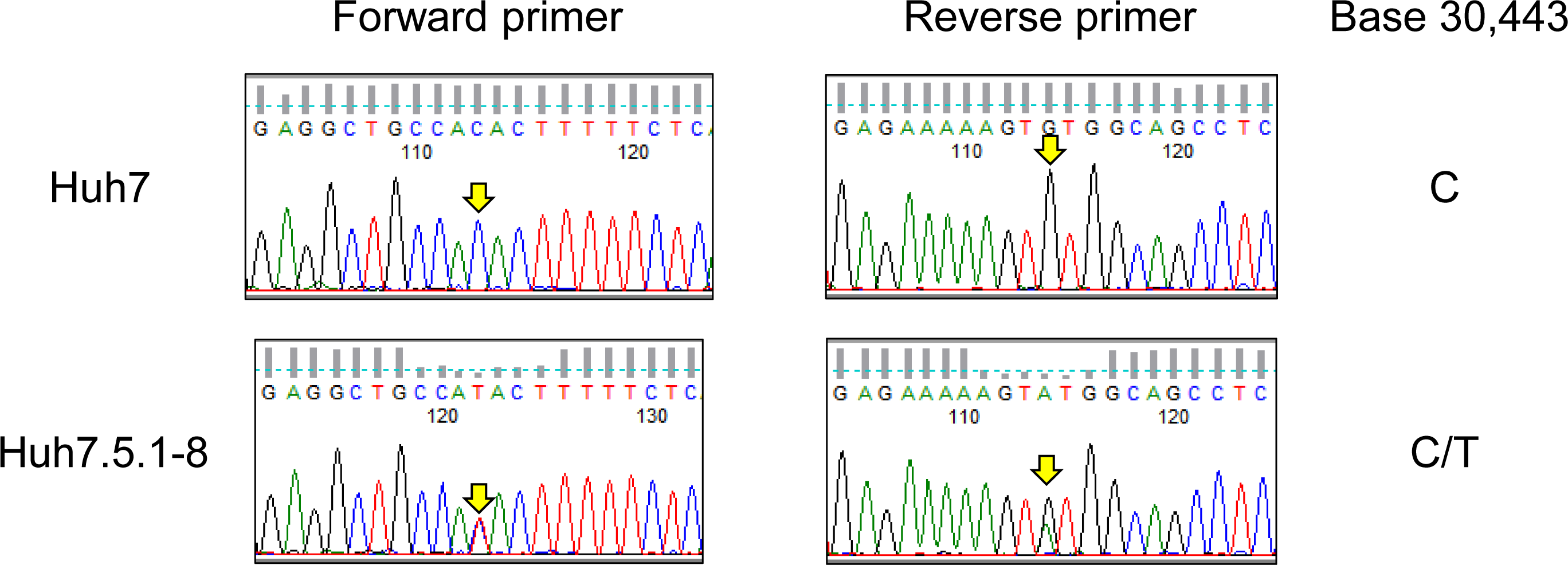
Sequence chromatograms of *DDX58* in Huh7 and Huh7.5.1-8. The genomic regions harboring the exon 2 of *DDX58* were amplified from Huh7 and Huh7.5.1-8 DNA samples using PCR and sequenced by the Sanger method. The arrows indicate a base 30,443 on NG_046918.1, and C-to-T transition at this position causes a T55I substitution of the DDX58 amino acid sequence.

In summary, the whole-genome sequencing of Huh7 and Huh7.5.1-8 provided various genetic characteristics of these cell lines, and some of them are, presumably, applicable to an in-house test for authentication of the Huh7 cell lineage. In addition, 53 genes were found to carry missense or LoF mutations specifically found in the Huh7 cell lineage. Among them, 8 genes contained mutations observed only in Huh7.5.1-8 or mutations with higher frequency in Huh7.5.1-8. These mutations might be relevant to the phenotypic differences between Huh7 and Huh7.5.1-8 and might also serve as genetic markers to distinguish Huh7.5.1-8 cells from the ancestral Huh7 cells.

## Acknowledgments

We thank the JCRB Cell Bank for providing DNA samples of Huh7 and Huh7.5.1-8. This work was supported by the Japan Society for the Promotion of Science, KAKENHI (grant 17H04003 to K.H. and grant 18H05511 to N.O.) and AMED-CREST (grant JP18gm0910005j0004 to K.H.).

## Author Contributions

Study design: T.E., M.F., K.H., and N.O., performing experiments: T.Y. and K.Sai: data analysis, M.K., K. Sat, and N.O., writing manuscript: M.K., T.Y., K.Sai., K.H., and N.O.

## Conflict of Interest

The authors declare no competing interests.

## Notes

#### Summary of Updates

The position of mutations in SPP1 was wrongly assigned. The mutation was changed from K200T to K241T in the main text and Table 2.

## References

Asselah, T., Bièche, I., Laurendeau, I., Paradis, V., Vidaud, D., Degott, C., et al. (2005). Liver Gene Expression Signature of Mild Fibrosis in Patients With Chronic Hepatitis C. Gastroenterology 129(6), 2064–2075. doi: https://doi.org/10.1053/j.gastro.2005.09.010.

Benjamini, Y., and Hochberg, Y. (1995). Controlling the False Discovery Rate: A Practical and Powerful Approach to Multiple Testing. Journal of the Royal Statistical Society. Series B (Methodological) 57(1), 289–300.

Blight, K.J., McKeating, J.A., and Rice, C.M. (2002). Highly Permissive Cell Lines for Subgenomic and Genomic Hepatitis C Virus RNA Replication. Journal of Virology 76(24), 13001–13014. doi: 10.1128/jvi.76.24.13001-13014.2002.

Bolger, A.M., Lohse, M., and Usadel, B. (2014). Trimmomatic: a flexible trimmer for Illumina sequence data. Bioinformatics 30(15), 2114–2120. doi: 10.1093/bioinformatics/btu170.

Bukh, J. (2016). The history of hepatitis C virus (HCV): Basic research reveals unique features in phylogeny, evolution and the viral life cycle with new perspectives for epidemic control. Journal of Hepatology 65(1, Supplement), S2–S21. doi: https://doi.org/10.1016/j.jhep.2016.07.035.

Chen, X., Schulz-Trieglaff, O., Shaw, R., Barnes, B., Schlesinger, F., Källberg, M., et al. (2016). Manta: rapid detection of structural variants and indels for germline and cancer sequencing applications. Bioinformatics 32(8), 1220–1222. doi: 10.1093/bioinformatics/btv710.

Choi, Steve S., Claridge, Lee C., Jhaveri, R., Swiderska-Syn, M., Clark, P., Suzuki, A., et al. (2014). Osteopontin is up-regulated in chronic hepatitis C and is associated with cellular permissiveness for hepatitis C virus replication. Clinical Science 126(12), 845–855. doi: 10.1042/cs20130473.

Cingolani, P., Platts, A., Wang, L.L., Coon, M., Nguyen, T., Wang, L., et al. (2012). A program for annotating and predicting the effects of single nucleotide polymorphisms, SnpEff. Fly 6(2), 80–92. doi: 10.4161/fly.19695.

Eckner, R., Ewen, M.E., Newsome, D., Gerdes, M., DeCaprio, J.A., Lawrence, J.B., et al. (1994). Molecular cloning and functional analysis of the adenovirus E1A-associated 300-kD protein (p300) reveals a protein with properties of a transcriptional adaptor. Genes & Development 8(8), 869–884. doi: 10.1101/gad.8.8.869.

Edgar, R.C. (2004). MUSCLE: multiple sequence alignment with high accuracy and high throughput. Nucleic Acids Research 32(5), 1792–1797. doi: 10.1093/nar/gkh340.

Fahmy, A.M., and Labonté, P. (2017). The autophagy elongation complex (ATG5-12/16L1) positively regulates HCV replication and is required for wild-type membranous web formation. Scientific Reports 7, 40351. doi: 10.1038/srep40351 https://www.nature.com/articles/srep40351#supplementary-information.

Feigelstock, D.A., Mihalik, K.B., Kaplan, G., and Feinstone, S.M. (2010). Increased susceptibility of Huh7 cells to HCV replication does not require mutations in RIG-I. Virology Journal 7(1), 44. doi: 10.1186/1743-422x-7-44.

Fujimoto, A., Furuta, M., Shiraishi, Y., Gotoh, K., Kawakami, Y., Arihiro, K., et al. (2015). Whole-genome mutational landscape of liver cancers displaying biliary phenotype reveals hepatitis impact and molecular diversity. Nature Communications 6, 6120. doi: 10.1038/ncomms7120 https://www.nature.com/articles/ncomms7120#supplementary-information.

Ghandi, M., Huang, F.W., Jané-Valbuena, J., Kryukov, G.V., Lo, C.C., McDonald, E.R., et al. (2019). Next-generation characterization of the Cancer Cell Line Encyclopedia. Nature 569(7757), 503–508. doi: 10.1038/s41586-019-1186-3.

Gilbert, D.A., Reid, Y.A., Gail, M.H., Pee, D., White, C., Hay, R.J., et al. (1990). Application of DNA fingerprints for cell-line individualization. American journal of human genetics 47(3), 499–514.

Iqbal, J., Sarkar-Dutta, M., McRae, S., Ramachandran, A., Kumar, B., and Waris, G. (2018). Osteopontin Regulates Hepatitis C Virus (HCV) Replication and Assembly by Interacting with HCV Proteins and Lipid Droplets and by Binding to Receptors αVβ3 and CD44. Journal of Virology 92(13), e02116–02117. doi: 10.1128/jvi.02116-17.

Jiao, X., Sherman, B.T., Huang, D.W., Stephens, R., Baseler, M.W., Lane, H.C., et al. (2012). DAVID-WS: a stateful web service to facilitate gene/protein list analysis. Bioinformatics 28(13), 1805–1806. doi: 10.1093/bioinformatics/bts251.

Johansson, E., and MacNeill, S.A. (2010). The eukaryotic replicative DNA polymerases take shape. Trends in Biochemical Sciences 35(6), 339–347. doi: https://doi.org/10.1016/j.tibs.2010.01.004.

Kahles, F., Findeisen, H.M., and Bruemmer, D. (2014). Osteopontin: A novel regulator at the cross roads of inflammation, obesity and diabetes. Molecular Metabolism 3(4), 384–393. doi: https://doi.org/10.1016/j.molmet.2014.03.004.

Kasai, F., Hirayama, N., Ozawa, M., Satoh, M., and Kohara, A. (2018). HuH-7 reference genome profile: complex karyotype composed of massive loss of heterozygosity. Human Cell. doi: 10.1007/s13577-018-0212-3.

Kim, D., Langmead, B., and Salzberg, S.L. (2015). HISAT: a fast spliced aligner with low memory requirements. Nature Methods 12, 357. doi: 10.1038/nmeth.3317 https://www.nature.com/articles/nmeth.3317#supplementary-information.

Knyaz, C., Stecher, G., Li, M., Kumar, S., and Tamura, K. (2018). MEGA X: Molecular Evolutionary Genetics Analysis across Computing Platforms. Molecular Biology and Evolution 35(6), 1547–1549. doi: 10.1093/molbev/msy096.

Koboldt, D.C., Zhang, Q., Larson, D.E., Shen, D., McLellan, M.D., Lin, L., et al. (2012). VarScan 2: Somatic mutation and copy number alteration discovery in cancer by exome sequencing. Genome Research 22(3), 568–576. doi: 10.1101/gr.129684.111.

Levine, B., Mizushima, N., and Virgin, H.W. (2011). Autophagy in immunity and inflammation. Nature 469, 323. doi: 10.1038/nature09782.

Li, H., and Durbin, R. (2009). Fast and accurate short read alignment with Burrows–Wheeler transform. Bioinformatics 25(14), 1754–1760. doi: 10.1093/bioinformatics/btp324.

Liu, G., Fan, X., Tang, M., Chen, R., Wang, H., Jia, R., et al. (2016). Osteopontin induces autophagy to promote chemo-resistance in human hepatocellular carcinoma cells. Cancer Letters 383(2), 171–182. doi: https://doi.org/10.1016/j.canlet.2016.09.033.

McKenna, A., Hanna, M., Banks, E., Sivachenko, A., Cibulskis, K., Kernytsky, A., et al. (2010). The Genome Analysis Toolkit: A MapReduce framework for analyzing next-generation DNA sequencing data. Genome Research 20(9), 1297–1303. doi: 10.1101/gr.107524.110.

Mills, R.E., Luttig, C.T., Larkins, C.E., Beauchamp, A., Tsui, C., Pittard, W.S., et al. (2006). An initial map of insertion and deletion (INDEL) variation in the human genome. Genome research 16(9), 1182–1190. doi: 10.1101/gr.4565806.

Minocherhomji, S., Ying, S., Bjerregaard, V.A., Bursomanno, S., Aleliunaite, A., Wu, W., et al. (2015). Replication stress activates DNA repair synthesis in mitosis. Nature 528, 286. doi: 10.1038/nature16139 https://www.nature.com/articles/nature16139#supplementary-information.

Nakabayashi, H., Taketa, K., Miyano, K., Yamane, T., and Sato, J. (1982). Growth of Human Hepatoma Cell Lines with Differentiated Functions in Chemically Defined Medium. Cancer Research 42(9), 3858–3863.

Ogryzko, V.V., Schiltz, R.L., Russanova, V., Howard, B.H., and Nakatani, Y. (1996). The Transcriptional Coactivators p300 and CBP Are Histone Acetyltransferases. Cell 87(5), 953–959. doi: https://doi.org/10.1016/S0092-8674(00)82001-2.

Oltval, Z.N., Milliman, C.L., and Korsmeyer, S.J. (1993). Bcl-2 heterodimerizes in vivo with a conserved homolog, Bax, that accelerates programed cell death. Cell 74(4), 609–619. doi: https://doi.org/10.1016/0092-8674(93)90509-O.

Osada, N., Kohara, A., Yamaji, T., Hirayama, N., Kasai, F., Sekizuka, T., et al. (2014). The Genome Landscape of the African Green Monkey Kidney-Derived Vero Cell Line. DNA Research 21(6), 673–683. doi: 10.1093/dnares/dsu029.

Perz, J.F., Armstrong, G.L., Farrington, L.A., Hutin, Y.J.F., and Bell, B.P. (2006). The contributions of hepatitis B virus and hepatitis C virus infections to cirrhosis and primary liver cancer worldwide. Journal of Hepatology 45(4), 529–538. doi: https://doi.org/10.1016/j.jhep.2006.05.013.

Petracca, R., Falugi, F., Galli, G., Norais, N., Rosa, D., Campagnoli, S., et al. (2000). Structure-function analysis of hepatitis C virus envelope-CD81 binding. Journal of virology 74(10), 4824–4830.

Pietrocola, F., Lachkar, S., Enot, D.P., Niso-Santano, M., Bravo-San Pedro, J.M., Sica, V., et al. (2014). Spermidine induces autophagy by inhibiting the acetyltransferase EP300. Cell Death And Differentiation 22, 509. doi: 10.1038/cdd.2014.215 https://www.nature.com/articles/cdd2014215#supplementary-information.

Sakuma, C., Sekizuka, T., Kuroda, M., Kasai, F., Saito, K., Ikeda, M., et al. (2018). Novel endogenous simian retroviral integrations in Vero cells: implications for quality control of a human vaccine cell substrate. Scientific Reports 8(1), 644. doi: 10.1038/s41598-017-18934-2.

Scheel, T.K.H., and Rice, C.M. (2013). Understanding the hepatitis C virus life cycle paves the way for highly effective therapies. Nature Medicine 19, 837. doi: 10.1038/nm.3248.

Shirasago, Y., Sekizuka, T., Saito, K., Suzuki, T., Wakita, T., Hanada, K., et al. (2015). Isolation and Characterization of an Huh.7.5.1-Derived Cell Clone Highly Permissive to Hepatitis C Virus. Japanese Journal of Infectious Diseases 68(2), 81–88. doi: 10.7883/yoken.JJID.2014.231.

Sumpter, R., Loo, Y.-M., Foy, E., Li, K., Yoneyama, M., Fujita, T., et al. (2005). Regulating Intracellular Antiviral Defense and Permissiveness to Hepatitis C Virus RNA Replication through a Cellular RNA Helicase, RIG-I. Journal of Virology 79(5), 2689–2699. doi: 10.1128/jvi.79.5.2689-2699.2005.

Szklarczyk, D., Franceschini, A., Wyder, S., Forslund, K., Heller, D., Huerta-Cepas, J., et al. (2015). STRING v10: protein–protein interaction networks, integrated over the tree of life. Nucleic Acids Research 43(D1), D447–D452. doi: 10.1093/nar/gku1003.

Tay, N., Chan, S.-H., and Ren, E.-C. (1990). Detection of integrated hepatitis B virus DNA in hepatocellular carcinoma cell lines by nonradioactive in situ hybridization. Journal of Medical Virology 30(4), 266–271. doi: doi:10.1002/jmv.1890300407.

Toshiyuki, M., and Reed, J.C. (1995). Tumor suppressor p53 is a direct transcriptional activator of the human bax gene. Cell 80(2), 293–299. doi: https://doi.org/10.1016/0092-8674(95)90412-3.

Wakita, T., Pietschmann, T., Kato, T., Date, T., Miyamoto, M., Zhao, Z., et al. (2005). Production of infectious hepatitis C virus in tissue culture from a cloned viral genome. Nature Medicine 11, 791. doi: 10.1038/nm1268 https://www.nature.com/articles/nm1268#supplementary-information.

Wang, J., Wang, W., Li, R., Li, Y., Tian, G., Goodman, L., et al. (2008). The diploid genome sequence of an Asian individual. Nature 456(7218), 60–65. doi: 10.1038/nature07484.

Wang, K., Yuen, S.T., Xu, J., Lee, S.P., Yan, H.H.N., Shi, S.T., et al. (2014). Whole-genome sequencing and comprehensive molecular profiling identify new driver mutations in gastric cancer. Nature Genetics 46, 573. doi: 10.1038/ng.2983 https://www.nature.com/articles/ng.2983#supplementary-information.

Wang, Q., Jia, P., and Zhao, Z. (2015). VERSE: a novel approach to detect virus integration in host genomes through reference genome customization. Genome Medicine 7(1), 2. doi: 10.1186/s13073-015-0126-6.

Yoneyama, M., Kikuchi, M., Natsukawa, T., Shinobu, N., Imaizumi, T., Miyagishi, M., et al. (2004). The RNA helicase RIG-I has an essential function in double-stranded RNA-induced innate antiviral responses. Nature Immunology 5, 730. doi: 10.1038/ni1087 https://www.nature.com/articles/ni1087#supplementary-information.

Zhong, J., Gastaminza, P., Cheng, G., Kapadia, S., Kato, T., Burton, D.R., et al. (2005). Robust hepatitis C virus infection <em>in vitro</em>. Proceedings of the National Academy of Sciences of the United States of America 102(26), 9294–9299. doi: 10.1073/pnas.0503596102.

