## Supplementary Table 1 for "Identification of Characteristic Genomic Markers in Human Hepatoma Huh7 and Huh7.5.1-8 Cell Lines"

Suppementary Table 1. PCR primer sequences

| ID | Primer s | Primer as1 | Primer as2 |
|---|---|---|---|
| DL1 | CATAGTTGACTTCACTGATGTGGAGC | CCATTATCAGTATCCCTCACCAGAG | CACTTGTCCTGCTGGCATTTGTATG |
| DL2 | GGGATGAAATAGCAGAGAGTGAGTTG | CATGGTATCCTCTGGACACATACTG | GATTCTTGGGACACTCAACATGTAGC |
| DL3 | GACAACTGTAGTCAATATAACATGGTACGTG | CTATCCAGATTCTCTCAAGCTTTCTCC | CTTTGCCAGGCACCTATTCATTAGC |
| DS1 | CAGTTGTCCACCTATGTGTGGG | CAGAAATATGAAGCTGCTGGAAAATATGATCC | NA |
| DS2 | CTAGCAACGGATCCAGTTCCAAG | GTCTGAAACCTTCTTCTCTCAACTCATC | NA |
| DS3 | GCTTGTGACATTAGAAACAAGATGGAGTG | CATCCTTGTTTCTAGTTACTGCCTTCC | NA |
