## Supplementary Table 2 for "Identification of Characteristic Genomic Markers in Human Hepatoma Huh7 and Huh7.5.1-8 Cell Lines"

Suppementary Table 2. Result of PCR amplifications for newly-identified deletions

| ID | Chromosome | Start position | End position | Δ Length | PCR result | Genes included |
|---|---|---|---|---|---|---|
| DL1 | 5 | 108094005 | 108356656 | 262652 | Huh7 and Huh7.5.1-8 specific Δ | *FBXL17,* exon |
| DL5 | 1 | 242396813 | 242421446 | 24634 | Huh7 and Huh7.5.1-8 specific Δ | *PLD5* , intron |
| DL6 | 5 | 97461126 | 97468551 | 7426 | Huh7 and Huh7.5.1-8 specific Δ but heterozygous in Huh-7 | |
| DS5 | 3 | 55290941 | 55291887 | 947 | Huh7.5.1-8 specific Δ | |
| DS6 | X | 120811663 | 120812521 | 859 | Huh7 and Huh7.5.1-8 specific Δ | |
| DS7 | X | 46301831 | 46302475 | 645 | Huh7.5.1-8 specific Δ | |
